# The Influence of PTH 1-34 on the Osteogenic Characteristics of Adipose and Bone Marrow Derived Stem Cells from Juvenile and Ovarectomized Rats

**DOI:** 10.1101/325134

**Authors:** Liza Osagie-Clouard, Anita Sanghani-Kerai, Melanie Coathup, catherine Pendegrass, Timothy Briggs, Gordon Blunn

## Abstract

Background: Mesenchymal Stem Cells (MSCs) are of growing interest in terms of bone regeneration; the majority of preclinical trials utilise bone marrow derived stem cells (bMSCs), though this is not without isolation and expansion difficulties

Objective: We compare the characteristics of bone marrow and adipose derived cells from juvenile, adult and ovarectomized rats; also assessing the effect of hPTH 1-34, on their osteogenic potential.

Methods: cells were isolated from the adipose and bone marrow of juvenile, adult and previously ovarectomized wistar rats. Cells were characterised with flowcytometery, proliferation assays, osteogenic and adipogenic differentiation, and migration to SDF-1. Experiments were repeated with and without co-culturing with 50nMol of intermittent PTH 1-34.

Results: The juvenile and adult MSCs demonstrated significantly increased differentiation into bone and fat and superior migration towards SDF-1 than ovarectomized groups, this was the case for adipose and bone marrow derived cells equally. PTH increased parameters of osteogenic differentiation and migration to SDF-1, this was significant for all cell types, though had the most significant effect on cells derived from OVX animals. Bone marrow derived cells from all groups, showed increased mineralisation and migration to SDF-1 compared to adipose derived cells.

Conclusion: Juvenile MSCs showed significantly greater migration to SDF-1 and showed greater osteogenic and adipogenic differentiation compared to cells from osteopenic rats, this was true for bone marrow and adipose derived cells. The addition of PTH, increased the osteogenic characteristics and migration of all cells, and further illustrates the possible clinical utility of both PTH and MSCs from various sources in bone regenerative therapies

## Introduction

Multiple influences alter the osteogenic capabilities of undifferentiated mesenchymal stem cells (MSCs); studies have compared the functional differences between cells from adolescent and aged animals, with a smaller body of work exploring the role of source osteoporosis on cell activity (1–4). Resultantly, the use of cells, be they minimally manipulated or enhanced with pharmacological adjuncts, may be key to orthopaedic therapies particularly in aged and osteoporotic populations; a patient cohort growing in size and worsening the musculoskeletal burden.

Post-menopausal oestrogen deficiency leads to an uncoupling of the bone remodelling cycle, where upregulated osteoclast activity is matched only with aberrant osteoblast activity-resulting in bone resorption (5,6). As such, type I osteoporosis is synonymous not only with reduced bone strength and thus increased fracture risk, but also is characterised by a reduction in bone mass and altered trabecular microarchitecture hence bone fragility (7). In clinical applications utilizing a tissue engineering approach or cell therapy treatments, the majority of pre-clinical studies have used mesenchymal stem cells derived from bone marrow (bMSCs). Yet, clinically this is not without problems, including morbidity associated with obtaining cells from iliac crest puncture, low cell yield, and reduced potency following extensive passage (8). Zuk et al (9) initially described the use of cells obtained from subcutaneous adipose liposuction aspirate as a source rich in MSCs (AdMSCs). The initial population isolated contains haemopoietic cells, pericytes, and adipose cells-all of which are plastic adherent; though after in vitro culturing, cells exhibit a homogeneous phenotype that fulfill the criteria for MSCs. Moreover, unlike periosteal cells or cells obtained from myogenic sources, adipose tissue is readily available, harvesting carries very limited morbidity, and cell yield is much greater than found from other sources (10,11). Reports on the osteogenic capacity of AdMSCs compared to bone marrow derived counterparts are contradictory; BMSC characteristics are thought to be affected by age, unlike AdMSCs where cells are thought to retain all characteristics regardless of age of source. Cell yield is also a fundamental difference between the sources, BMSCs yield 6 × 10^6^ nucleated cells per mL of aspirate, with a maximum of 0.01% being MSCs. Whereas, 2 × 10^6^ cells can be isolated from 1 gm adipose tissue, and 10% are stem like (12,13). As such, investigations into the combined effect of age and ovarectomy on the osteogenic efficacy of these cells is pivotal to fully determine their possible eventual clinical utility.

In addition, to cell source and age of animal, pharmacological adjuncts can be used to modulate stem cell behavior. Studies, including those conducted on post-menopausal women, demonstrate a profound anabolic effect of parathyroid hormone 1-34 (PTH) (14,15). Moreover, in vitro data has shown PTH to mediate MSC fate, not only increasing MSC number, but also their preferential osteogenic differentiation over adipogenesis (16). Interestingly, these findings have all been reported in cells derived from bone marrow, with very little data on the effect of PTH on adipose derived MSCs.

The ability for intermittent PTH (iPTH) to mobilise/increase migration of cells from the haemopoietic niche, is particularly significant in the context of sites of increased bone turnover (fractures and peri-implant). Cells are initially “mobilised” from their niche into the circulation, “home” across the tissue endothelium and mature into active cell types-eventually “modulating” the local environment. The SDF-1/CXCR4 axis has been found to be an important regulator of stem cell migration. Stromal derived factor-1/CXC1L2 (SDF-1) is produced by a multitude of tissue types including fracture endosteum and in its active form is bound to the CXCR4 receptor found on MSCs. Granero-Molto et al (17) demonstrated that dynamic stem cell migration to the fracture site in a stabilised tibial osteotomy model was CXCR4 dependent. The clinical significance of the SDF-1/CXCR4 axis has further been alluded to, whereby the overexpression of CXCR4 on mesenchymal stem cells led to significant increases in bone mineral density; thus, having implications in the treatment of osteoporosis (18). Jung et al demonstrated increased osteoblast expression of SDF-1 following iPTH administration and thus upregulation of the stem cell homing axis SDF-1/CXCR4, alluding to significant implications for endochondral repair and increased bone volume fraction (19)

As such, although comparisons have been made between the osteogenic potential of different stem cell sources; a very limited body of work compares these differences across juvenile, adult and ovarectomized animals, nor does this work elucidate their capacity to migrate to SDF-1. Moreover, no study has explored this in the context of PTH administration. We hypothesise that mesenchymal stem cells isolated from bone marrow aspirates and adipose tissue of juvenile female Wistar rats will have greater tridiffrentiation, migration and proliferation than cells isolated from adult and ovarectomized rats, and that co-culture with intermittent PTH will upregulate the osteogenic characteristics of both adipose and bone marrow derived cells.

## Methods

### Isolation, culture and expansion of MSCs

Female Wistar rats were used throughout this study, animals were classed as "juvenile" at 2-4 weeks or "adult” at 6-9 months, one group of animals (“OVX”) were supplied immediately post ovarectomy (all animals supplied by Charles River Laboratories, Harlow, Essex). These animals were housed for 16 weeks in pairs and osteopenia confirmed by assessing their femoral, lumbar 3rd and 4^th^ vertebrae and humeri mineral density with pQ-CT compared to age matched non-ovarectomized controls; a reduction of 22% in bone mineral density was confirmed, as such our model was one of osteopenia rather than osteoporosis.

Husbandry conditions for all animals used throughout the thesis were standardised, whereby animals were housed in pairs in cages without restriction to ambulation on a standard rodent diet. Cages included adequate bedding and unrestricted access to fluids, along with objects for animal enrichment

#### Bone Marrow Cell Isolation

Following euthanasia, all animals were processed within 60minutes of euthanasia to maintain cell viability. Within a laminar flow hood, dissected femora were washed twice with phosphate buffered saline (PBS, ThermoFisher, and Hemel Hempstead, UK) to remove remaining external debris. Using sterile forceps, ends were transected at the diaphsyeal-metaphyseal junction leaving a diaphyseal portion of varying lengths between 30-40mm; the medullary canal was flushed three times using a 23 gauge needle with 5mls of Dulbecco’s Modified Eagle Medium high glucose (DMEM) was flushed through the shaft, with the aspirate collected and cultured in DMEM, 20% fetal calf serum and 1% Penicillin Streptomycin (“standard media”).

#### Adipose derived stem cell Isolation

Adipose derived stem cells (AdMSC) were isolated from the abdominal subcutaneous fat, avoiding perinephric and visceral fat. Under aseptic conditions, adipose samples were washed three times with PBS following the removal of other soft tissues (muscle, tendon, ligaments) and weighed. Subsequently, specimens were minced with sterile scissors, In a 20ml universal tube, 8mls of warmed 0.1% collagenase-type II (Sigma-Aldrich) was added to the adipose tissue and agitated in a 37°C water bath for 60minutes. Samples were then centrifuged at 5000rpm for 5minutes, forming a gelatinous pellet. The supernatant was aspirated and discarded, leaving the remaining pellet in the tube, to which a further 5mls of standard media was added, followed by repeat centrifuge.

Following this, the supernatant was discarded, and the pellet resuspended in 5mls of fresh standard media, the suspension was then cultured in standard media, and again used for experimental procedures at passage 3-4 (40).

### Characterisation of Stem Cells-Tridiffrentiation

Osteogenic: All experiments were repeated in triplicate on cell cultures from 3 different animals per group. 30,000 confluent cells were cultured in supplemented standard media-termed herein as “osteogenic media” containing, 1×10^−7^M water soluble Dexamethasone (Sigma-Aldrich,UK), 1×10^−4^M Ascorbic acid (Sigma-Aldrich, Dorset, UK) and 1×10^−2^M Beta-glycerol phosphate (Sigma-Aldrich, Dorset, UK). Cells were cultured in these conditions for 21 days

#### Alizarin Red Staining

30.0 Seeded cells were cultured in either standard or osteogenic media for 21 days in duplicate, from juvenile, adult and ovarectomized senile rats, N=3.

At days 7, 14 and 21 mineralisation was assessed by staining calcium deposits with Alizarin Red. Cells were fixed in formalin and stained with 100μl of alizarin red solution, the plates covered with foil and incubated at room temperature for 15minutes. The stain was then aspirated and the cells washed multiple times with PBS until the solution ran clear to remove nonspecific staining. Cells were then left in 100μl of PBS and images taken.

To quantify staining, 10% cetylpyridinium chloride (CPC) was added to 10mM of sodium phosphate to obtain a working solution of pH 7. Following imaging, PBS was discarded from wells, and 200μl of the CPC solution added to each well for 15minutes, agitated at room temperature and covered in foil. Absorbance was then read on a plate reader at wavelength 570nm (Tecan infinite Pro, Tecan trading, Switzerland).

A standard curve was made by serial dilution of alizarin red working solution in CPC and read again at 570nm.

#### Osteocalcin Staining

15,000 cells were seeded in 96 well plates for each group (N=3), the media aspirated and cells washed twice with PBS. They were then fixed in 10% formalin at room temperature for 10 minutes, and washed twice with cold PBS, cells were then incubated in blocking buffer (PBS, 0.1% tween20 and 1% bovine serum albumin) for 1hour at room temperature. Anti rat osteocalcin (Abcam, Cambridge UK) was diluted in the blocking buffer (1:1000), and the cells incubated in this solution overnight at 4C. Cells were then washed three times with PBS and incubated for 2hours in the dark in Goat Anti-Mouse IgG H&L (Abcam, Cambridge UK) also diluted in the blocking buffer (1:100). After dilution, the secondary was removed and the cells again washed with PBS. Cells were then counterstained by incubating in DAPI (Abcam, Cambridge UK) for 20minutes, again diluted in the blocking buffer (1:1000), rinsed with PBS and imaged under a fluorescent microscope.

#### Alkaline phosphatase (ALP) activity

Osteoblast differentiation was measured using an alkaline phosphatase assay (N=3 per group), cells were seeded in 48-well plates at a concentration of 4500 cells/cm^2^, and cultured in either standard or osteogenic media. ALP data was collected at days 3,7,14 and 21 and repeated in duplicate at each time point.

50μl of each sample (repeated in duplicate) was combined in a 96-well nunc microplate reader with 50μl of p-Nitrophenol phosphate (Sigma-Aldrich, Dorset, UK), and agitated at 37C in the dark for 30minutes. Plates were then read at a fluorescence of 405nm and measurements expressed in U/L.

Adipogenic: 30,000 confluent cells were cultured in supplemented standard media-termed herein as “adipogenic media”, consisting of 0.5mM isobultyl-1-methylxanthine (Sigma-Aldrich, Dorset, UK), 10ug/ml insulin (Sigma-Aldrich, Dorset, UK) and 1uMl dexamethasone (Sigma-Aldrich, Dorset UK).

#### Oil Red O staining

30,000 cells were cultured in adipogenic media for 21 at days 7, 14 and 21 lipid production was assessed by staining droplets with Oil Red O. to fixed cells 100ml of 60% isopropanol was added to each well and incubated for 15minutes at room temperature, cells were then washed twice with PBS and incubated in Oil Red O working solution for 15minutes at room temperature. After staining, cells were washed carefully with 60% isopropanol and analysed. To quantify staining, 200ul of 100% isopropanol was added to each well, and the plate agitated at room temperature for 15minutes. The supernatant was then collected and absorbance read on a plate reader at wavelength 510nm.

A standard curve was made by serial dilution of Oil Red O working solution in isopropanol and read again at 510nm

Cell morphology. Passage 2 and 3 cells from the three groups for both sources were cultured in standard media and their morphology was assessed by measuring their aspect ratio, whereby the ratio of the length of a cell to its width was calculated using ImageJ software.

### 2.2.3 Characterisation of stem cells-Flocytometery

A total of 100 000 cells from the bone marrow and adipose of young, adult control, and OVX rats were analysed for their surface expression of cluster of differentiation (CD) CD29, CD90, CD45,CD106, CD146 and CD34. The CD expression was compared with the isotype control. Cells were fixed in 4% formalin for 15 minutes at room temperature, washed with 0.5% bovine serum albumin (BSA), and stained with the conjugated primary antibody for one hour at room temperature in the dark. After one hour, the cells were washed with 0.5% BSA and analysed on flow cytometer (Cytoflex, Beckman Coulter, Brea, California).

### 2.2.3 Preparation of PTH

Human parathyroid hormone 1-34 (Teriparatide, Bachem, Hertfordshire UK) was dissolved in 4mM hydrochloric acid solution containing 0.1% bovine serum albumin to a stock concentration of 5mM, which was subsequently stored at-20C. Intermittent regimens involved the culture of cells in the PTH containing media for 6hours in every 48 hour cycle at 50nMol based on a preliminary study comparing dosing regimens and concentrations (20). We repeated the osteogenic differentiation of cells (N=3 for each group from both tissue sources) with the addition of iPTH-again assessing with ALP, alizarin red and osteocalcin immunocytochemistry.

#### The effect of PTH 1-34 on cell proliferation

10,000 cells derived from the fat and bone marrow of juvenile, adult, and OVX rats (n = 3 for each group) were incubated in DMEM, 20% fetal calf serum, 1% penicillin streptomycin with iPTH. At 3,7,10 and 14 days 10% Alamar Blue assay (AbD Serotec, Kidlington, United Kingdom) was added to the culture media for four hours; the resultant media was read at an excitation of 560 nm and an emission of 590 nm using a Tecan plate reader (Infinite Pro 200 Series, Tecan, Mannedof, Switzerland). The mean absorbance was determined from the triplicate samples. The absorbance was then normalized to the DNA assays and a comparison was made between the groups

#### The effect of PTH 1-34 on cell migration to SDF-1

At passage 3 cells were seeded at a concentration of 4500cells/cm^2^ in 48 well plates and cultured with either osteogenic media or osteogenic media supplemented with 50nM of PTH 1-34 for 21 days. After which cells were trypsinised and counted using a haemocytometer. 10,000 cells from each group were loaded in plain DMEM without supplements in the upper compartments of a Boyden chambers (Merck, Watford UK). The lower compartment of the chambers was filled with 100ng/ml SDF-1 in standard media, and incubated at 37°C, 5% CO2.

After 16 hours, the cells that migrated to the opposite side of the membrane were fixed in 10% formalin and stained with Toluidine blue.

Following fixation, the chambers were rested in 200μl of the toluidine solution for 3minutes. Subsequently, wells were washed three times with distilled water, after which wells were analysed under the microscope, where cells that had migrated were counted by selecting six random fields at x20 magnification and calculating the percentage average number of cells. For the control, both the top and bottom of the chamber were filled with standard media with no SDF1 and the cells were loaded in the upper chamber as described.

Statistical Analysis: Values are expressed as the mean ± SD. Experiments were performed at least in triplicate. Data was found to be non parametric following Shapiro Wilkinson testing, and as such Mann-Whitney U tests were used for analysis in Graphpad software (GraphPad Software, Inc., San Diego, CA)

## 2.3 Results

Expression of CD markers: There was no significant difference in CD marker expression by cells obtained from any group, regardless of tissue source. Mean CD marker expression is outlined in Table 1

**Table 1.** Cluster differentiation marker expression showed no significant difference between groups.

|  | Adipose | Bone marrow |
| --- | --- | --- |
| **Juvenile** | CD29 (mean 95.8%)<br>CD90 (mean 91.6%)<br>CD 106 (mean 86%)<br>CD 146 (mean 90.2%)<br>CD34 (mean 2.7% )<br>CD45 (mean 9.1%) | CD29 (mean 95.1%)<br>CD90 (mean 96.6%)<br>CD 106 (mean 88%)<br>CD 146 (mean 91%)<br>CD34 (mean 2.6%)<br>CD45 (mean 10.9%) |
| **Adult** | CD29 (mean 90.9%)<br>CD90 (mean 94.7%)<br>CD 106 (mean 85%)<br>CD 146 (mean 89.6%)<br>CD34 (mean 3.6%)<br>CD45 (mean 12.9%) | CD29 (mean 91.1%)<br>CD90 (mean 97%)<br>CD106 (mean 85%)<br>CD146 (mean 89%)<br>CD34 (mean 3.8%)<br>CD45 (mean 9%) |
| **OVX** | CD29 (mean 97.2%)<br>CD90 (mean 91.9%)<br>CD 106 (mean 87.1%)<br>CD 146 (mean 89.7%)<br>CD34 (mean 1.7%)<br>CD45 (mean 9.7%) | CD29 (mean 99.1%)<br>CD90 (mean 94.4%)<br>CD 106 (mean 87%)<br>CD 146 (mean 90%)<br>CD34 (mean 1.7%)<br>CD45 (mean 11.1%) |

Cell morphology: Both adipose derived and bone marrow derived MSCs from juvenile rats demonstrated a tight spindle like morphology, with no significant difference in mean aspect ratios (bMSC 18.66 and aMSC 19.1). The mean ratios in adult cells were significantly smaller (bMSC 4.99 and aMSC 5.31), though again there was no difference between different tissue sources. MSCs from ovx rats had the smallest aspect ratio compared to the other cell types (bMSC 2.25 AdMSC 1.80) (Figures 1).

**Figure 1:**
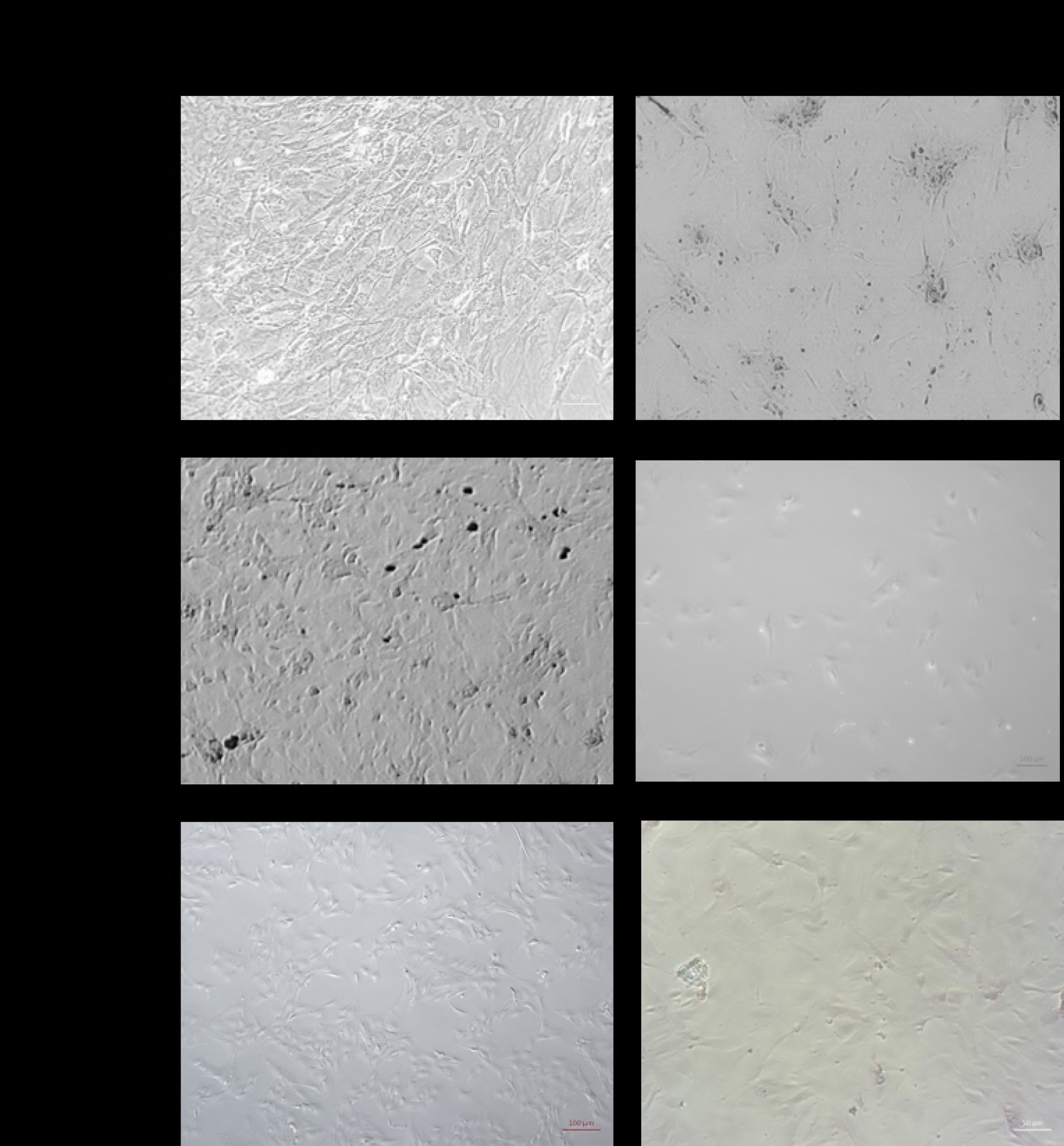
Images of MSCs derived from the bone marrow and adipose tissue of a) juvenile, b) adult, c) OVX ratsProliferation: No effect on cell proliferation normalised against DNA was seen secondary to age or ovarectomy, nor were any differences noted between the different tissue sources.

### Tridiffrentiation of cells

Cells from all sources were able to tridiffrentiation with the addition of specific culture media.

Osteogenic: Mineralization of the extracellular matrix increased in all groups over the 21 day experimental period. At day 7, juvenile bone marrow derived cells produced significantly more calcium phosphate than OVX cells (p = 0.004); this trend continued over the 21day period. There was no difference between calcium phosphate deposition from juvenile and adult derived bMSCs at any time point, this was also the case for adipose derived cells. This trend was repeated in adipose derived cells whereby juvenile cells at all timepoints had significantly greater mineralisation than OVX cells (P <0.07) when comparing tissue source bone marrow derived cells deposited significantly more calcium phosphate then adipose derived cells, this difference was most profound for ovx cells (p=0.4 juvenile, p=0.4 adult, p=0.05 ovx) (Figures 2,3)

On the addition of iPTH, cells showed a significant increase in alizarin red staining compared to untreated groups at all time points for bone marrow derived cells. This effect was noted to be most profound on OVX cells that showed a nearly 2fold increase on calcium phosphate deposition compared to untreated cells at day 21(p<0.05) This effect was also seen in adipose derived OVX cells

Bone marrow derived cells demonstrated the most significant reaction to 50noml of iPTH compared to adipose derived cells by day 21 (ovx p=0.05, juvenile p=0.4, adult p=0.4). (Figure 2,3,4)

There was large variability in the ALP expression from day three to day 14 across all groups. no difference between ALP expression from cells derived from adipose or bone marrow was seen, but as with calcium phosphate deposition, juvenile and adult cells expressed significantly more ALP than OVX cells at day 14 when production peaked for cells from both tissue sources (p<0.04). iPTH led to increased ALP production peaking at day 14 for all cell types compared to untreated cells (p=0.05); this effected adipose derived and bone marrow cells to the same degree, with no difference in the magnitude of the effect independent of age or ovarectomy.

**Figure 2:**
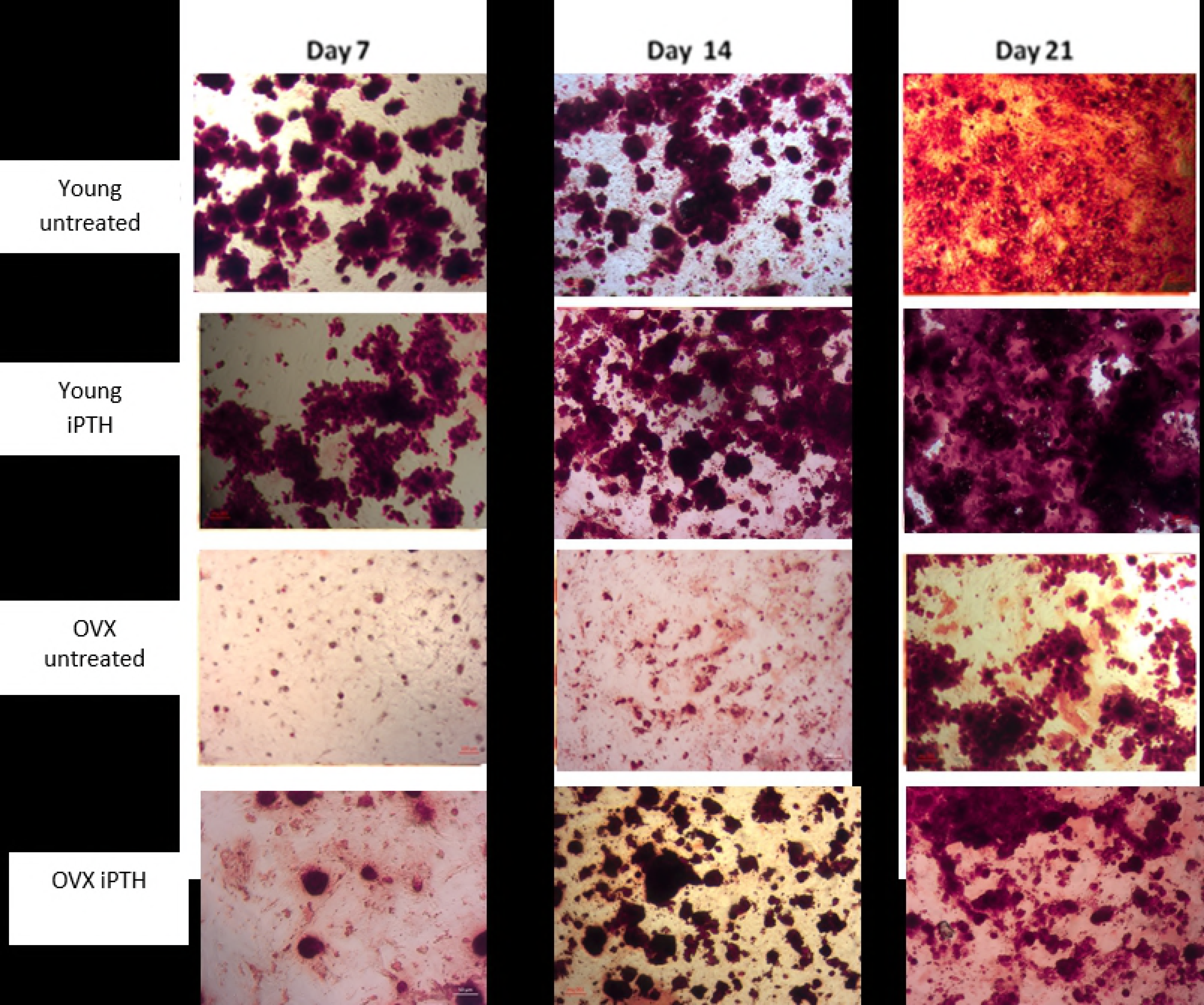
Effect of 50nM iPTH on alizarin red staining of young and OVX adipose derived cells PTH

**Figure 3:**
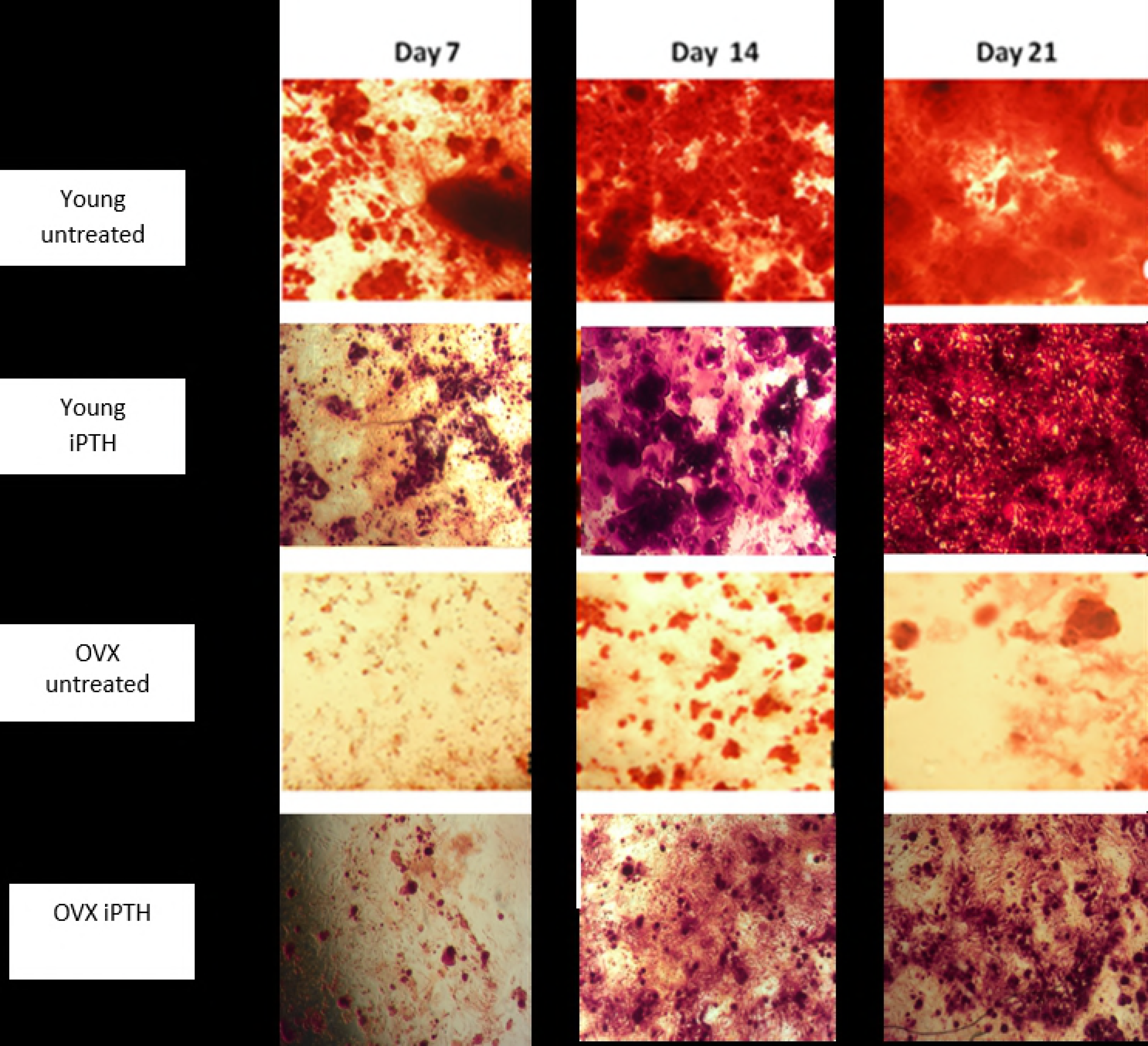
Effect of 50nMol iPTH on alizarin red staining of young and OVX bone marrow derived cells.

**Figure 4:**
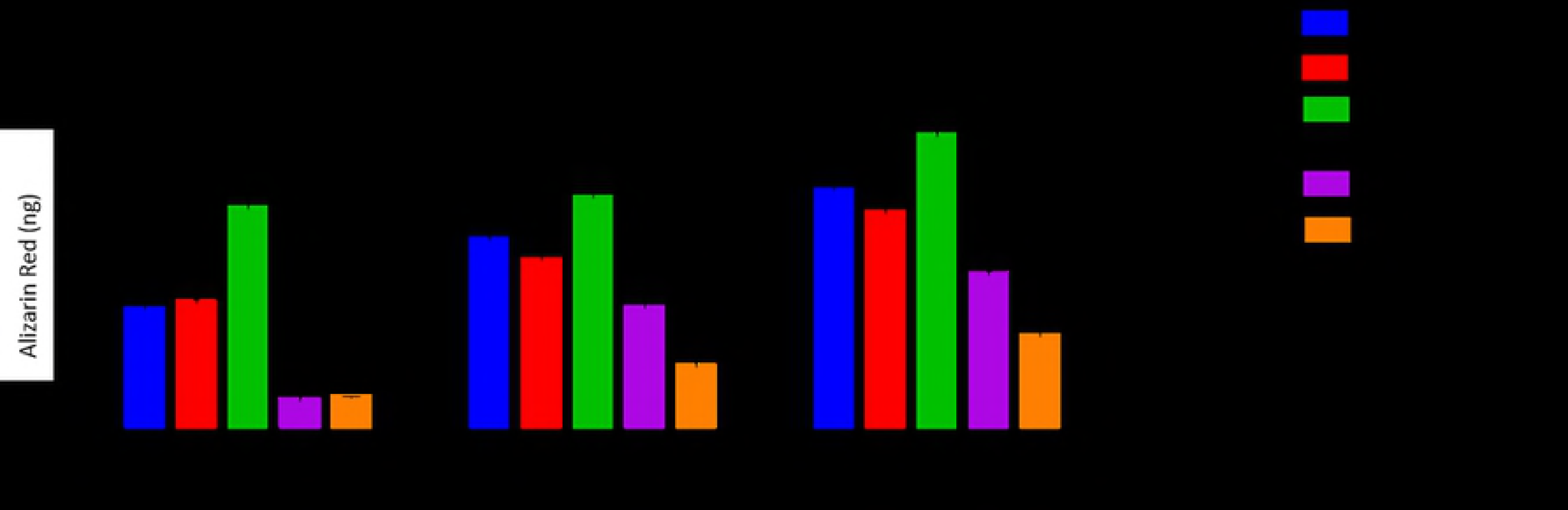
Graph of alizarin red staining. Juvenile bone marrow cells, produced significantly more calcium phosphate at days 7 and 14 than OVX cells *p<0.04. This was also evident in adipose derived cells **p<0.07. Bone marrow derived cells produced significantly more stained calcium phosphate than adipose derived cells at every time point *** p<0.05 (for juvenile cell comparison)

Cells cultured in intermittent PTH showed greater staining for osteocalcin at day 7 compared to cells cultured in osteogenic media alone; this was true of cells derived from both juvenile, adult and ovarectomized rats from bone marrow and adipose tissue. Though cells from ovarectomized rats treated and untreated, exhibited less staining than their juvenile and adult counterparts. No discernible difference could be seen between staining for adipose or bone marrow derived cells for osteocalcin staining (Figure 5).

**Figure 5:**
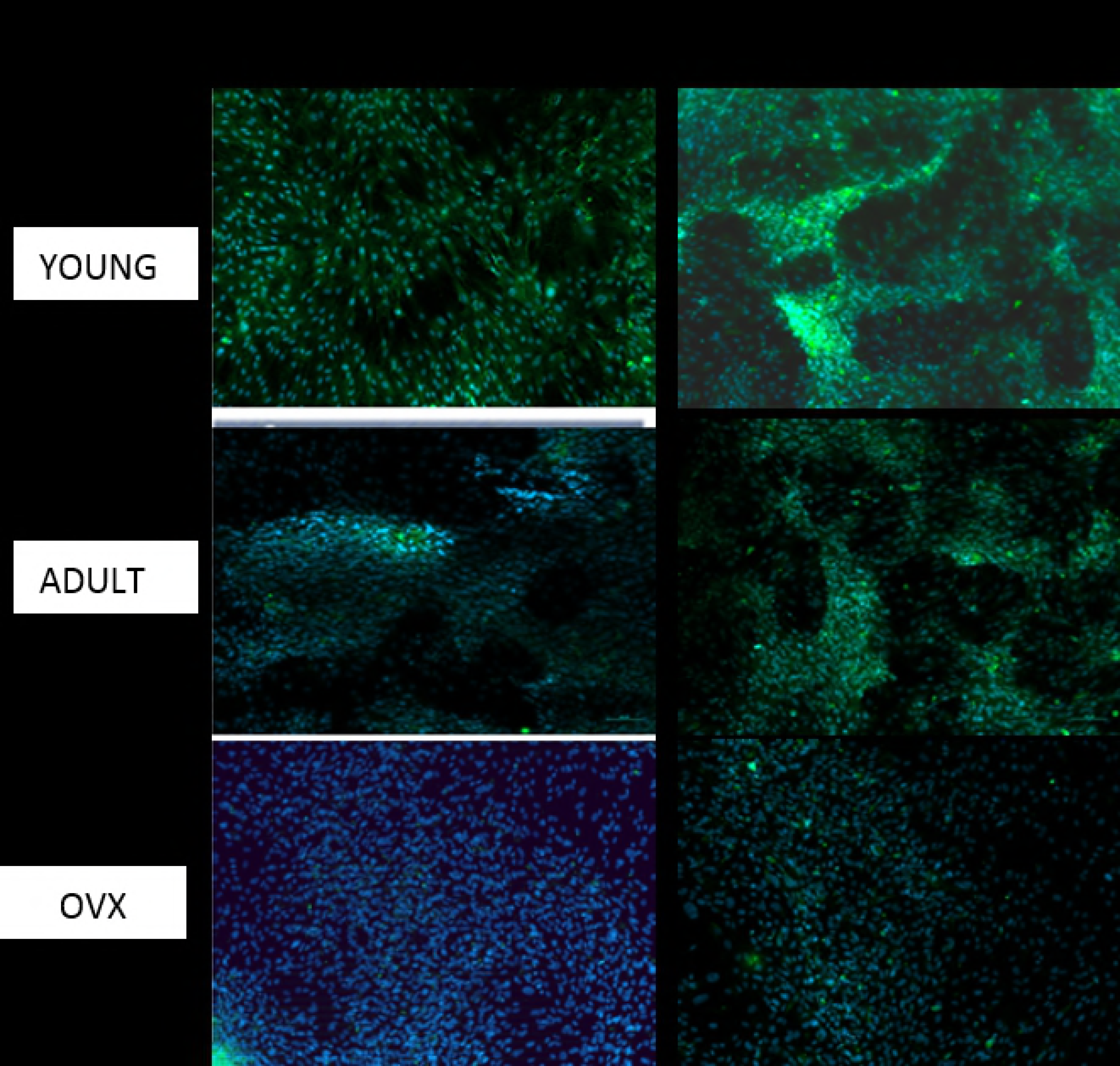
Osteocalcin immunocytochemistry at day 7 for bone marrow derived cells, comparing images from cells cultured with and without 50nM of intermittent PTH 1-34

Adipogenic: At days 14 and 21, adipocytic differentiation was significantly greater in MSCs isolated from juvenile animals compared with adult control and OVX groups for both tissue sources. MSCs from young rats accumulated significantly greater amounts of lipid from day seven compared with the other two groups of cells. The rate of lipid accumulated from day 7 onwards was greater in cells isolated from juvenile rats.

Additionally, lipid droplet accumulation significantly accelerated from day 14 to day 21 for juvenile bMSCs, while bMSCs from adult and ovX rats showed no increase in lipid accumulation after day 7. Cells isolated from adipose tissue regardless of donor age or whether they were derived from OVX rats continued to show increased lipid formation over the 21 day period, though juvenile cells were always more productive than the other groups.

When comparing bone marrow to adipose derived cells, there was no difference in adipogenic differentiation, but the reduction in micro droplet formation was more significant between adipose juvenile and ovx cells, than seen with bone marrow derived cells (p<0.06)

Cell Migration to SDF-1: In bone marrow derived cells, the migration of SDF-1 in the juvenile cell group was significantly greater than all groups (p=0.05) and was nearly twice as high as the migration of OVX cells (p<0.05). In adipose derived cells, the migration of MSCs from young rats was significantly less than with bone marrow derived cells though the pattern of these cells migrating more than cells from OVX animals was continued (P<0.07) (Figure 8). On the addition of continuous PTH, no significant difference was found in migration, yet on adding 50nmol of iPTH, all cell types demonstrated increased migration compared to their untreated counterparts and this treatment affected adipose and bone marrow derived cells equally (figure 6, 7a, b, 8 a,b).

**Figure 6:**
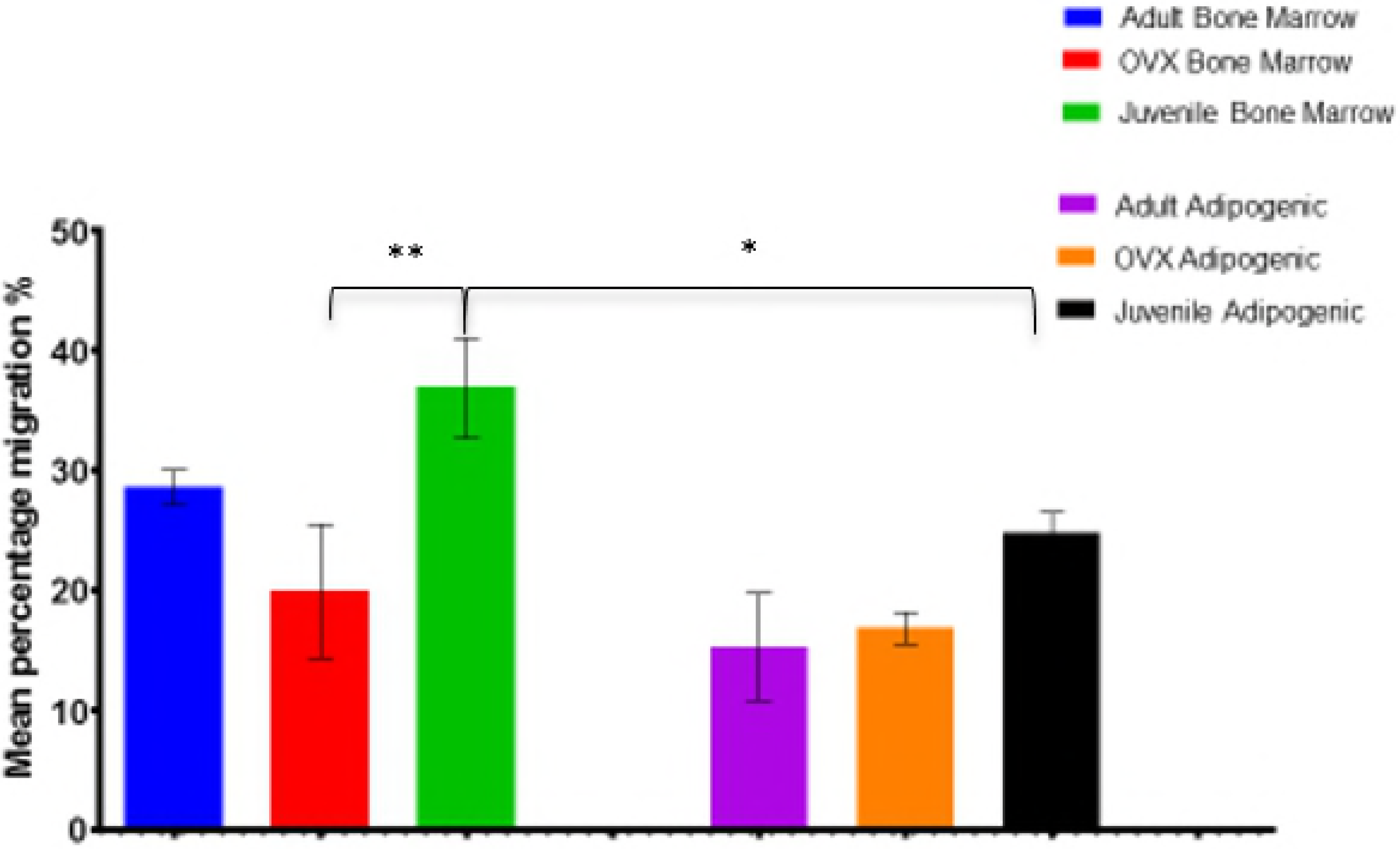
Graph demonstrating cellular migration to SDF-1. Juvenile cells migrated significantly more than OVX derived cells in both groups p=0.05** BM, p=0.07 Ad). While bone marrow cells migrated more than adipose derived cells.

**Figure 7a,b:**
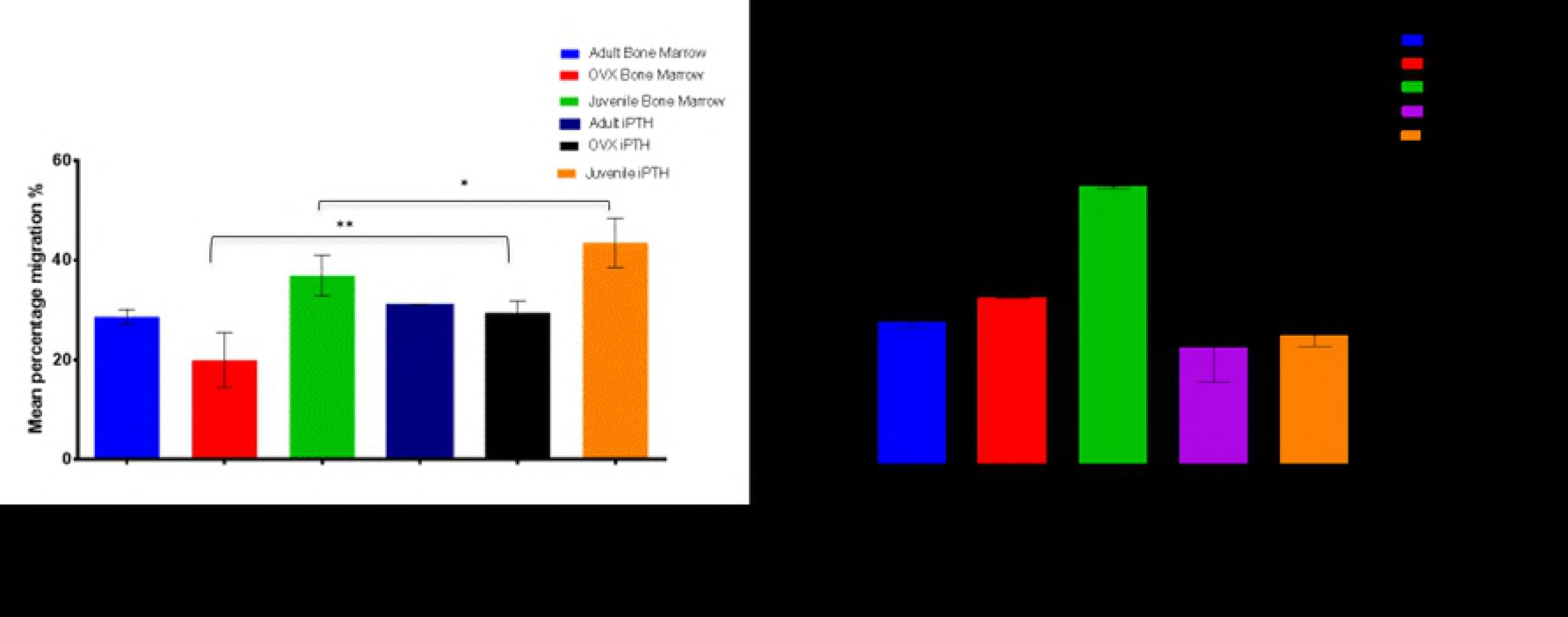
a-Graph demonstrating cellular migration to SDF-1 in bone marrow derived cells with 50nMol iPTH. Juvenile cells showed increased migration with iPTH (p=0.05*), as did OVX cells (p=0.04**) b-Graph demonstrating cellular migration to SDF-1 in adipose derived cells with 50nMol iPTH. Juvenile cells showed increased migration with iPTH (p=0.4*), as did OVX cells (p=0.5**)

**Figure 8a,b:**
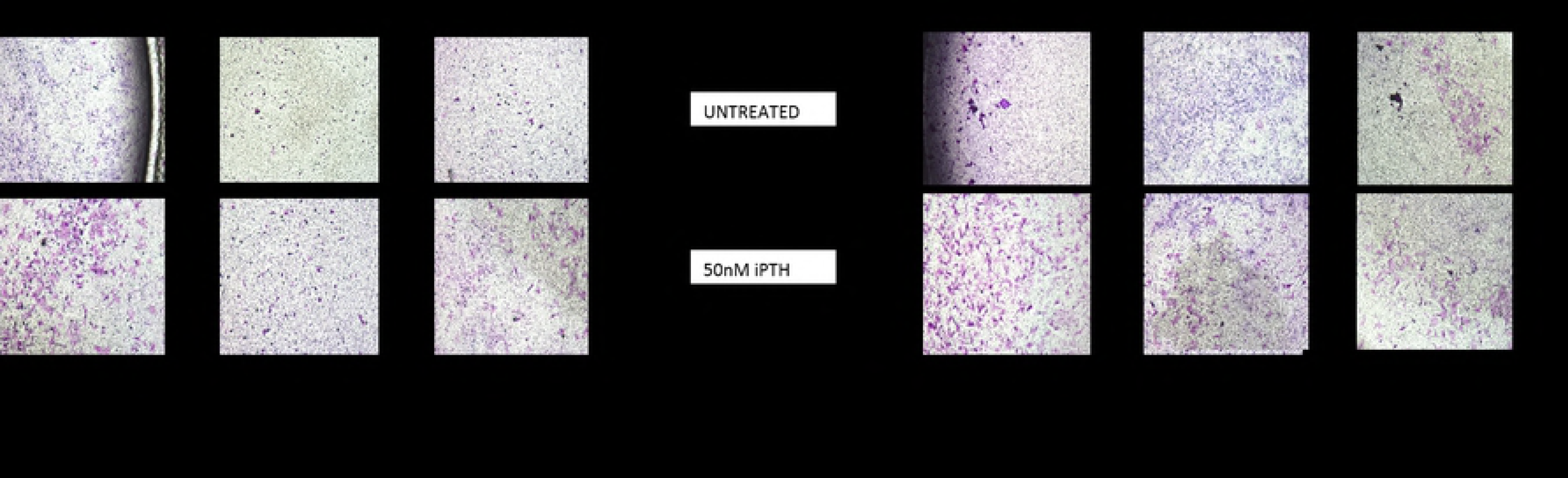
a-Images of bone marrow derived cells in Boyden chambers migrated to SDF-1, with and without iPTH 1-34 treatment b-Images of adipose derived cells in Boyden chambers migrated to SDF-1, with and without iPTH 1-34 treatment

## 2.4 Discussion

This study examines the capacity of adipose and bone marrow derived cells, to differentiate into adipocytes osteoblasts and chondrocytes and determined the effects of PTH 1-34 dosing regimens on cellular osteogenic characteristics and migratory capacity. we found that adipose derived cells demonstrated poorer calcium phosphate deposition, osteocalcin expression, and migration along the CXCR4/SDF-1 axis compared to bone marrow derived cells; and that these differences was more profound for cells derived from ovarectomized animals rather than their juvenile counterparts. We also found differences between cells isolated from juvenile and ovarectomized animals for each cell source, with a reduction in osteogenic and migrative characteristics in OVX derived cells although these cells showed similar proliferative capacity and CD marker expression. Finally, this work demonstrated increased osteogenic capabilities of young and OVX cells from both tissue sources, when dosed with pulsatile PTH 1-also resulting in increased migration of all cells, though had no effect on cellular proliferation. Interestingly this effect was often greater with OVX cells, and on cells obtained from bone marrow rather than adipose tissue.

My data showed no difference in CD marker expression from the cells independent of source or age or OVX status of animal. Other studies have demonstrated a higher expression of CD34 from adipose derived cells at early culture (11), this cluster differentiation marker is important for cell-to-cell adhesion, as well as cell extracellular matrix deposition. Similarly, several reports suggest higher CD106 and CD146 from bone marrow derived cells compared to adipose derived cells. My findings were contrary to this, whereby all samples were negative for CD34, and there was equal expression of CD106 and CD146. In my study this finding may be explained by the use of the cells at passage 3 particularly as previous work has demonstrated a reduction in CD34 expression at later passage (11). The lack of difference between OVX and juvenile derived cells and CD markers, is in keeping with other work (21), that being said, the marker heterogeneity in study methodologies means only a limited inference can be made between the expression of CD markers, and the in vivo/in vitro activity of cells and as such, the value of marker expression cannot be taken in isolation. Similarly, the value of morphology is limited in isolation, though we found no difference morphologically between adipose or bone marrow derived cells, both demonstrating the same spindle like phenotype from juvenile populations, and both moved morphologically to a more flattened phenotype from aged and ovx animals.

Moerman et al (22) reported changes in gene expression of aged rats compared to juvenile counterparts, with an upregulation in proliferator-activated receptor-gamma (PPAR-γ), but a down-regulation in BMP-2 and TGF-B in older rats, which would go some way in explaining the propensity of aged cells to typically differentiate preferentially down the adipogenic lineage as seen and the increased adiposity seen in the marrow of both aged and osteoporotic rats compared to their juvenile counterparts. Asumda and Chase demonstrated reduced osteogenic and adipogenic differentiation ability of MSCs from old rats compared to MSCs from young rats (23). However, Singh et al. (24) found no observable difference in osteogenic and adipogenic differentiation between cells from young and old mice. Likewise, Beane et al (25) showed no differences in alkaline phosphatase expression as well as alizarin red staining between bone marrow MSCs from young and old rabbits but they found that age affected the adipogenic differentiation of the same cells and also led to reduced adipogenic differentiation in adipose derived cells, findings we mirrored. My results for bone marrow derived cells, showed a marked reduction in the osteogenic and adipogenic differentiation between juvenile and OVX cells. The reason for the disparate findings between the studies is likely to be multifactorial. There are clear differences in the method of culture and the “osteogenic media” used to induce differentiation. Moreover, although differences were seldom seen between adult and juvenile cells, due to the time taken for osteopenia to develop, the ovx animals we used were older than all other groups-thus there would be benefit in further elucidating results by using a truly aged matched population.

Interestingly in our study, we found no significant difference in ALP expression with AdMSCs compared to BMSCs from both OVX and juvenile cells. This was not consistent with our other markers for osteogenic activity, which were significantly higher with BMSC derived cells. Chen et al (26) and Wu et al (27) compared the osteogenic potential of juvenile and aged AdMSCs, finding no difference in functional characteristics, though there was a discernible change in morphology and telomere length. Indeed, we found this to be true when cells isolated from juvenile and adult rats cells were compared, but when derived from ovx animals, there was a significant difference in all measured osteogenic outputs compared to juvenile derived cells, the reasons for which remain unclear though the cellular cytokine profile following ovarectomy is known to be significantly altered compared to age matched counterparts.

A fundamental advantage of using adipose rather than bone marrow derived cells in bone regeneration, is based on the evidence, that the proliferative and osteogenic capacity of adipose derived cells is not effected by age. In the present study, there was no difference in proliferation between any of MSCs independent of age, OVX or source. This is in contrast to other studies assessing bone marrow derived cells, whereby, MSCs from older rats have significantly lower proliferation compared to MSCs from young rats (28,29). Beane and co-workers (25) looked at MSCs from young and old rabbits derived from the bone marrow, muscle and fat, where the cells from bone marrow demonstrated a reduction in proliferation with age whereas cells from the other two sources did not. Georgen (30) found that MSCs from OVX rats had a lower proliferation rate than their control counterparts and concluded that the low proliferation rate would correlate with reduced self-renewal capacity which might cause a gradual depletion of MSC sources in the bone marrow of OVX animals.

The stimulation, proliferation and differentiation of bone marrow derived osteoprogenitor cells by PTH 1-34 has been well documented in the literature (31). The anabolic window is based on findings of intermittent dosing regimens, Contrary to the larger body of literature we did not find an increase in proliferation of cells from any group after PTH dosing, this is contrary to work where proliferative increases with shorter exposure cycles to PTH (30minutes-1hour), thought to be due to selective upregulation of the cAMP/PKA pathways (32).

When comparing the effects of intermittent PTH on the osteogenic capacity of cells derived from adipose and bone marrow, we found significant differences. Although PTH affected all OVX cells, this change was most significant with cells derived from bone marrow; where cell mediated mineralisation at day 7 for OVX and young bone marrow cells had increased 2.1 and 1.9 fold compared to comparative cells from adipose tissue, which had increased 1.6 and 1.5 times respectively.

The non-collagenous calcium binding protein, osteocalcin, is regarded as a late marker of osteogenic differentiation, immunohistochemistry showed greater expression from bone marrow derived young cells compared to OVX cells, while intermittent PTH led to increases in all groups. Similarly, young adipose derived cells had greater expression than OVX cells, with a similar effect of PTH. Although bone marrow MSC osteogenic characteristics remained superior to those of adipose derived cells, in both groups the production of osteocalcin was significantly enhanced on the addition of iPTH. These findings were similar to that of Zhang et al (33) and Park et al (34) who used mechanical and electrical stimulation respectively, and although BMSCs remained superior, they did see marked improvement in the activity of AdMSCs.

These experiments demonstrated MSCs from OVX rats have a lower in vitro migration compared to MSCs from juvenile and adult rats from both cell sources. That adipose derived cells had poorer migrative capacity than those from bone marrow. On the addition of intermittent PTH, all cells increased migration.

SDF-1 is a chemokine receptor for CXCR4 and the SDF-1/CXCR4 biological axis plays an important role in the migration of stem cells and the wound repair of tissues and organs. The impaired migration capacity of MSCs from 4 months post-ovariectomy rats may be due to their low expression of CXCR4 and hence this could explain the impaired bone formation in osteoporotic patients as these cells have a reduced capacity to migrate to the site of bone loss. SDF1 is produced in the periosteum of injured bone and encourages endochondral bone repair by recruiting mesenchymal stem cells to the site of injury. Therefore, mobilization of osteoblastic progenitors to the bone surface is an important step in osteoblast maturation and formation of mineralised tissue. Very little work has explored the effect of PTH on adipose derived cells; we found these cells also reacted to PTH in a similar way to bone marrow derived cells with the percentage increase in migration was greatest in OVX cells-compared to untreated cells.

The findings of this chapter confirm the ability to obtain stem cells from the adipose and bone marrow of Wistar rats; supporting the notion that although stem cells from both adipose and bone marrow remain active, their osteogenic differentiation ability is affected by ovarectomy therefore impairing their regenerative and differentiation capacity.

Interestingly in vivo studies have yielded mixed results on the efficacy of AdMSCs in bone formation. Nieymeyer et al (35) and Hayashi (36) showed significantly poorer fracture healing in sheep and rat defects respectively compared to the osteogenesis and full bone bridging achieved with implanted BMSCs. Indeed, implanted undifferentiated ADMSCs tended to differentiate into cells with an adipose like morphology and thus hindered healing. This is converse to Kang et al (37) and Stockmann et al (38) who in porcine and canine models respectively, found no difference in bone formation with BMSCs and ADMSCs. Yet, herein lies the difficulty in comparing in vivo studies; the heterogeneity of subcutaneous versus intra-abdominal fat, cell number, cell culture techniques and fracture model makes comparisons between studies difficult. What is clear is that cell yield and isolation of AdMSCs is greater and easier than with BMSCs. Hence in light of my results one wonders if PTH can enhance the osteogenic capabilities of AdMScs so that they are equivalent or better those of BMSCs which would significantly increase their clinical utility.

This study examines three very different variables, the effect of osteopenia/age, and the effect of cell tissue source and the role of PTH on all of these. I have found that migration of cells and osteogenic differentiation is reduced when derived from osteopenic animals, and that bone marrow derived cells have greater calcium phosphate deposition and osteocalcin expression. But importantly, we have also demonstrated that the addition of PTH will improve the activity of all cell types, Yukata et al (39) reported the reduced efficacy of periosteal stem cells in an aged osteopenic mouse tibial fracture model when treated with PTH compared to a juvenile model; our findings are converse to this and instead demonstrated increased sensitivity of OVX cells to PTH. This may have implications for clinical applications: If allogenic cells from younger patients are incompatible for use in the aged osteoporotic population, then can the addition of PTH improve the ability of OVX derived cells to migrate and differentiate in order to render them effective for bone regeneration?

A significant caveat to this work is that all experiments were carried out in 2-D, cell culture whereas in vivo chapters effects may be more complicated because of the fracture environment, the role of periosteal cells and post fracture genomic cascades which are not present in vivo.

Conflicts of Interest: No authors report a conflict of interest

## Acknowledgements/Funding

Works reported have been supported by a grant from the Rosetree’s Trust, and Gwen Fish Orthopaedic Trust

